# Discovery of additional ancient genome duplications in yeasts

**DOI:** 10.1101/2025.08.31.673279

**Authors:** Kyle T. David, Linda Horianopoulos, Carla Gonçalves, Jacob L. Steenwyk, Ana Pontes, Paula Gonçalves, Chris Todd Hittinger, Matt Pennell, Antonis Rokas

**Affiliations:** Department of Biological Sciences, Vanderbilt University, Nashville, TN 37235, USA; Evolutionary Studies Initiative, Vanderbilt University, Nashville, TN 37235, USA; Laboratory of Genetics, J. F. Crow Institute for the Study of Evolution, Center for Genomic Science Innovation, DOE Great Lakes Bioenergy Research Center, Wisconsin Energy Institute, University of Wisconsin-Madison, Madison, WI 53726, USA; Department of Food Science, University of Guelph, Guelph, ON, Canada, N1G 2W1; UCIBIO, i4HB, Departamento de Ciências da Vida, Faculdade de Ciências e Tecnologia, Universidade Nova de Lisboa, Caparica, Portugal; Howard Hughes Medical Institute and the Department of Molecular and Cell Biology, University of California, Berkeley, CA 94720, United States; Department of Computational Biology, Cornell University, Ithaca NY 14853 USA

**Keywords:** Whole Genome Duplication, Yeasts, Polyploidy, Convergent Evolution

## Abstract

Whole genome duplication (WGD) has had profound macroevolutionary impacts on diverse lineages^1,2^, preceding adaptive radiations in vertebrates^3–5^, teleost fish^6,7^, and angiosperms^8,9^. In contrast to the many known ancient WGDs in animals^10,11^ and especially plants^12–14^, we are aware of evidence for only four in fungi^15,16^. The oldest of these occurred ∼100 million years ago (mya) and is shared by ∼60 extant Saccharomycetales species^17,18^, including the baker’s yeast *Saccharomyces cerevisiae* (Fig. 1). Notably, this is the only known ancient WGD in the yeast subphylum Saccharomycotina. The dearth of ancient WGD events in fungi remains a mystery^15^. Some studies have suggested that fungal lineages that experience chromosome^19^ and genome^15^ duplication quickly go extinct, leaving no trace in the genomic record, while others contend that the lack of known WGD is due to an absence of data^15,16^. Under the second hypothesis, additional sampling and deeper sequencing of fungal genomes should lead to the discovery of more WGD events. Coupling hundreds of recently published genomes from nearly every described Saccharomycotina species with three additional long-read assemblies, we discovered three novel WGD events. While the functions of retained duplicate genes originating from these events are broad, they bear many similarities to the well-known WGD that occurred in the Saccharomycetales^17^. Our results suggest that WGD may be a more common evolutionary force in fungi than previously believed.

**Highlights:**

- Evidence for three whole genome duplications (WGDs) in Dipodascales yeasts
- Impacts of WGD are broad but bear many similarities to the known WGD in Saccharomycetales yeasts
- Duplicates with many protein-protein interactions are more likely to be retained over long timescales
- WGD in fungi is likely underreported

## Results and Discussion

### Evidence for three whole genome duplication events in the Dipodascales

To detect signatures of ancient WGD we first used a gene/species tree reconciliation algorithm to infer gene duplications across the Saccharomycotina phylogeny. High rates of gene duplication along specific lineages of the species tree can be indicative of WGD events and have been used to identify ancient WGDs in plants^20–22^, animals^23^, and fungi^24^. An initial analysis using a 400-species backbone phylogeny (Fig. S1) successfully recovered the known ancient WGD near the base of Saccharomycetales (labelled WGD1). This lineage possessed the second-highest rate of duplications, over 10x higher than the average internal lineage. However, the lineages with the first and third highest rates of duplication occurred in the Dipodascales, a clade separated from the Saccharomycetales by ∼300 million years (my) of evolution.

To improve the resolution of gene family evolution within both clades, we increased our sampling in Saccharomycetales (135 genomes) and Dipodascales (184 genomes) and performed additional gene/species tree reconciliation analysis. The lineage containing WGD1 was again successfully identified by a spike in duplication rate, experiencing 53.1 gene duplications / my (Fig. 1). Two internal spikes were also evident in Dipodascales. The most recent ancestral lineage to the genera *Dipodascus* and *Geotrichum* (spanning from 201-132 mya) had a duplication rate of 21.8 duplications / my, indicating that a WGD may have occurred along this lineage (labelled WGD2). Similarly, the most recent ancestral lineage to the species *Magnusiomyces tetraspermus* and *Saprochaete suaveolens* (spanning from 62-38 mya) had a duplication rate of 23.62 duplications / my (labelled WGD3) (Fig. 1). Additionally, the highest species-specific duplication rate by far belonged to *M. magnusii* with 130 gene duplications / my, 2.4x higher than any other Dipodascales species. *M. magnusii* also possesses the largest known Saccharomycotina genome (43.2Mb, 3.4x larger than *S. cerevisiae*), leading us to hypothesize another WGD (labelled WGD4, spanning from 27-0 mya) specific to this species alone.

**Figure 1.**
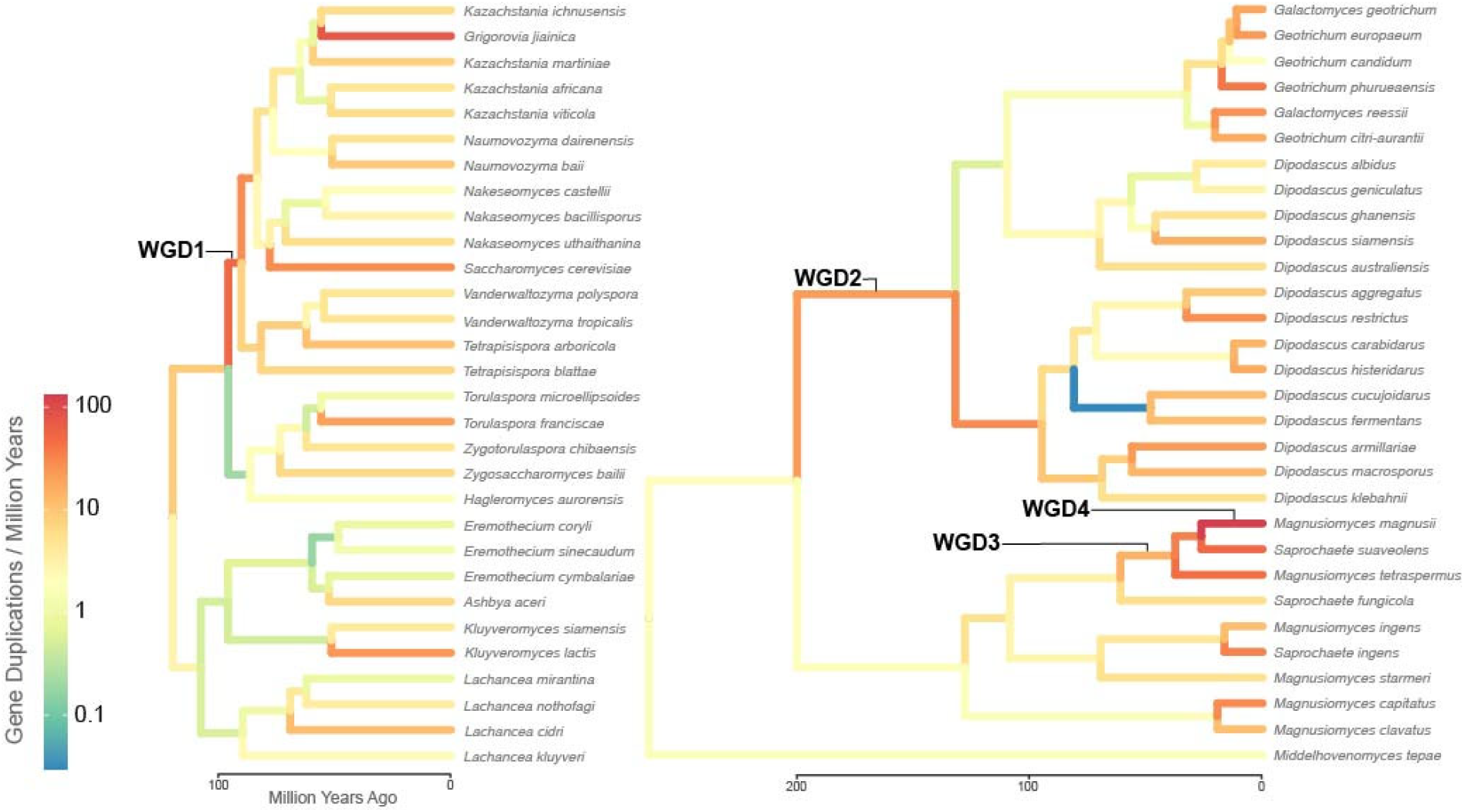
Three additional whole genome duplications in Saccharomycotina yeasts. Rates of gene duplication are mapped onto Saccharomycetales (left) and Dipodascales (right) phylogenies. Whole genome duplications indicated along the lineages in which they are predicted to occur. Trees have been pruned for visualization purposes.

While gene/species tree reconciliation approaches are useful in identifying putative WGDs, they are limited in their ability to distinguish true WGD from bouts of single or segmental gene duplication events^25^, or in definitively placing WGDs on specific lineages due to gene tree discordance and unbalanced paralog retention^24^. For example, descendant lineages of WGD2 and WGD3 also have elevated rates of gene duplication (Fig. 1).

A complementary approach to identify WGD involves the detection of colinear segments of paralogs (called multiplicons) within a genome^26,27^. Such analysis requires highly contiguous assemblies. Assemblies with fewer than 100 contigs were available for one post-WGD2 (*G. candidum*: 28 contigs) and one post-WGD3 (*Sap. suaveolens*: 12 contigs) species. To generate additional evidence for hypothesized WGDs and their placements on the yeast phylogeny we re-sequenced three additional species: *D. fermentans* for WGD2 (19 contigs), *M. tetraspermus* for WGD3 (33 contigs) and *M. magnusii* for WGD4 (71 contigs). These 5 genomes, along with 81 other highly contiguous Saccharomycotina genome assemblies (Data S1) were then searched for multiplicons (Fig. 2A).

**Figure 2.**
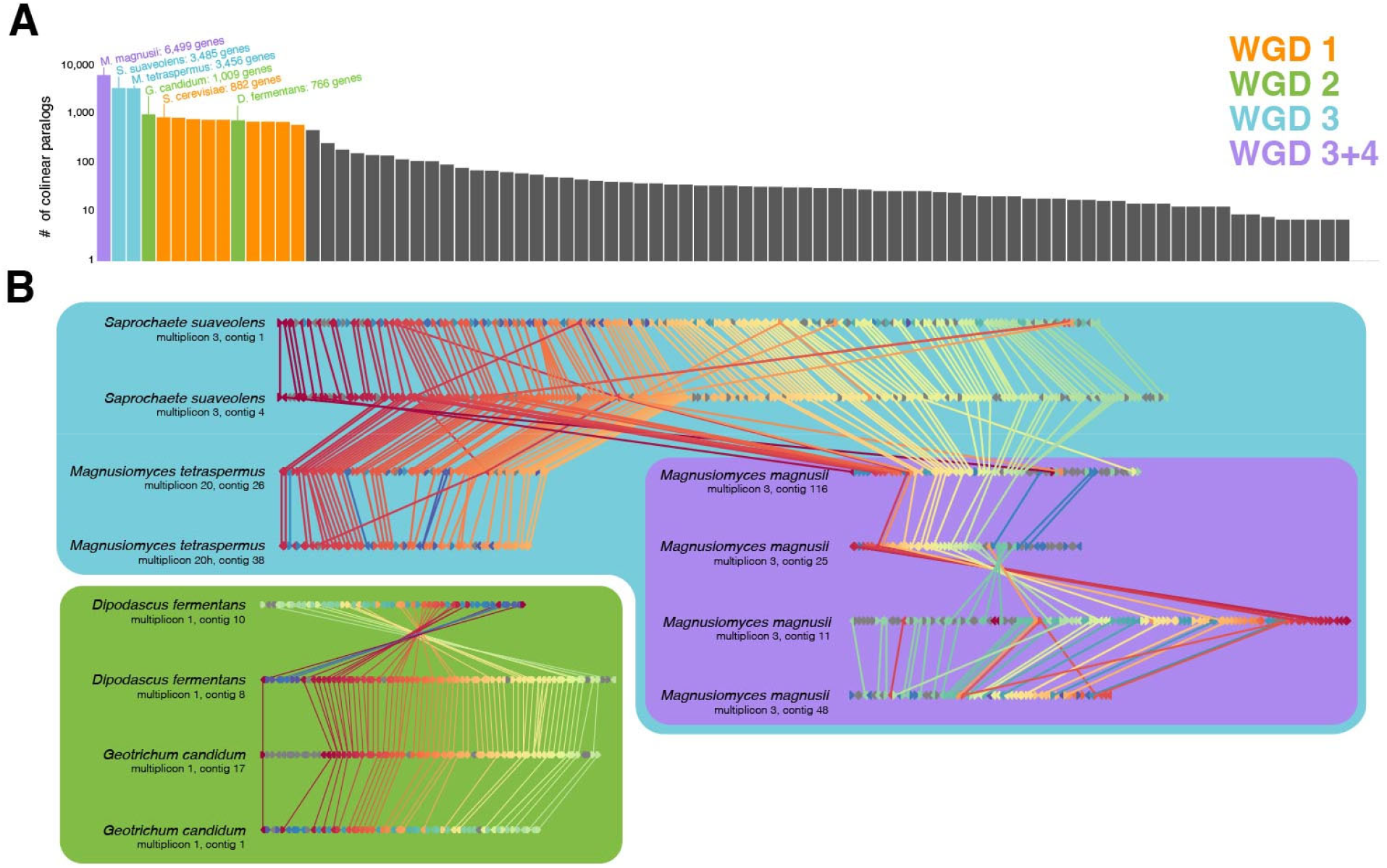
Paralog collinearity supports WGD events. A) Cumulative multiplicon size of each highly contiguous Saccharomycotina assembly, as measured by the number of unique genes. Genomes predicted to have undergone ancient WGD are highlighted. B) Gene order of select multiplicons. Colors represent unique orthogroups, demonstrating homology both within and between species. Orthogroups that only appear once are depicted in gray.

We found that genomes hypothesized to descend from ancient WGDs each contained numerous large multiplicons across scaffolds. Furthermore, multiplicons between *D. fermentans* and *G. candidum* had many orthogroups in common, as did *Sap. Suaveolens, M. tetraspermus*, and *M. magnusii* (Fig. 2B). However, multiplicons between post-WGD2 and post-WGD3 species (*D. fermentans* and *Sap. suaveolens*, for example) contained largely distinct orthogroups (Data S2), consistent with our inference that WGD2 and WGD3 represent two distinct events (Fig. 1). Interestingly, whereas multiplicons for most post-WGD species occurred in duplicates, in *M. magnusii*, they appear in tetraplicate, providing further evidence for WGD4 as an event specific to this species. WGD4 is also supported by examination of the cumulative multiplicon size in *M. magnusii*, which is 186.5% that of sister species *Sap. Suaveolens*.

Another common hallmark of ancient WGD is large increases in the number of chromosomes across lineages^28^. However, fungi possess small, loosely-packed chromosomes, which make karyotyping difficult^29^. Furthermore, Saccharomycotina yeasts have extremely diverse telomeres^30,31^ and centromeres^32,33^, which make chromosome number estimation challenging even for highly-contiguous assemblies. Despite these obstacles, chromosome count estimates from several Dipodascales, species exist, including one representative from each proposed WGD. For species not inferred to have undergone WGD, count estimates range from four^34^ to five^35,36^ chromosomes. This estimate increases to seven chromosomes for the post-WGD3 species *Sap. suaveolens*^37^ and eight^38^ or nine^39^ for the post-WGD2 species *G. candidum*. Finally, 13 chromosomes have been inferred from pulsed field gel electrophoresis for *M. magnusii*^40^, which is predicted to have undergone two rounds of WGD (both WGD3 and WGD4). Therefore, while limited chromosome count estimates in Dipodascales preclude a formal analysis, we interpret the available data to be consistent with our conclusions.

WGD can occur both within lineages (autopolyploidization) or between lineages via hybridization^24,41^ (allopolyploidization). WGD1 was originally hypothesized to be an autopolyploidization event due to the highly conserved gene order within descendent genomes^18^. However, a gene/species tree reconciliation analysis later recovered a lineage pre-dating WGD1 with a higher rate of gene duplications than the WGD1 lineage itself^24^. The authors posit this unusual finding could be explained by gene tree discordance produced through hybridization^24^, which suggests allopolyploidization. As our reconciliation analysis recovered no such peak in gene duplication rates preceding any of the novel WGD events (Fig. 1), we hypothesize that these events occurred either through autopolyploidization or through allopolyploidization between closely related parent species.

In contrast to more ancient WGD events, WGD4 may have occurred relatively recently and is apparently specific to just *M. magnusii*. Thus, this WGD may be of similar age to several young hybrids within Saccharomycotina species complexes that are known to vary in ploidy levels^42–47^. Though *M. magnusii* is not a known hybrid, further investigation is warranted to better elucidate the timing and mechanism of WGD4. For example, broader sampling within *M. magnusii* will help determine whether WGD4 is shared by the entire species, or specific to just the taxonomic type strain.

### Common consequences of genome duplication

WGD is often referred to as an engine of evolutionary innovation^2,4^. For example, it has been hypothesized that WGD1 facilitated aerobic glycolysis in several yeast species^48,49^. Also known as the “Crabtree/Warburg effect”, this is the biochemical process that empowers baking, brewing, and winemaking. However, others have noted that some species that have not undergone WGD1 are still capable of aerobic glycolysis to a lesser extent^50^. As this example illustrates, identifying cause-effect relationships from rare evolutionary events, such as WGD, is exceedingly difficult^51,52^. The discovery of additional WGD events in Saccharomycotina affords new opportunities for understanding how WGD events contribute to evolutionary innovation.

To test whether certain functional gene classes were more likely to be retained in duplicate following WGD, and if these classes were convergently shared across events, we ran enrichment analysis on retained orthologs for each event. Notably, WGD4 ohnologs were significantly enriched (adjusted p<0.05) for only five InterPro Gene Ontology^53^ categories (Data S3) and for none of the Kyoto Encyclopedia of Genes and Genomes^54^ (KEGG) pathways (Data S4). As 70% of all *M. magnusii* genes are ohnologs, we hypothesize that not enough time has passed since WGD4 for genes to become lost and patterns within retained copies to emerge. As discussed above, it is not yet known whether WGD4 represents a truly ancient WGD or a strain-specific allopolyploidization. Therefore, we turn our attention to the three older WGD1, WGD2, and WGD3 (WGD1,2,3 for short) for the remainder of this discussion.

The effects of WGD appear widespread. Ohnologs were significantly enriched (adjusted p<0.01) in at least six of the seven main KEGG pathway categories across WGD1,2,3 (Data S4). Despite the diversity of affected pathways, commonalities remain. Enrichment analysis of InterPro Gene Ontology annotations and KEGG pathways revealed three overarching functional themes in post-WGD genomes: metabolism, expression, and signaling (Fig. 3). We discuss each of these in turn below.

**Figure 3.**
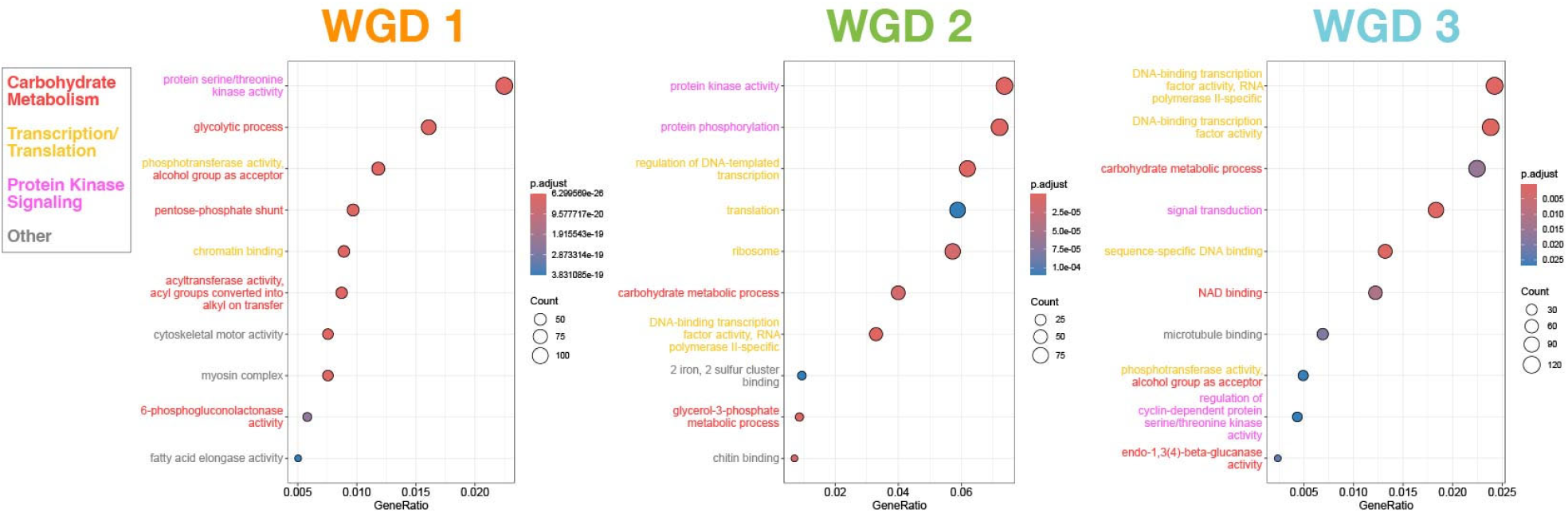
Shared gene ontology categories are preferentially retained following ancient WGD in different clades of Saccharomycotina yeasts. The ten most overrepresented gene ontology categories in ohnologs from WGD1, WGD2, and WGD3 are shown, colored by generic category.

### Metabolism

Within the KEGG carbohydrate metabolism subcategory, no specific pathway was shared between events. However, four related pathways were significantly enriched (adjusted p<0.01) in one of the three WGD events (Fig. 4). For example, WGD2 ohnologs were enriched in the propanoate metabolic pathway. Propanoate esters and other related compounds are responsible for the characteristic fruity scent of *G. candidum*^55^, a popular yeast in cheesemaking^56^ as well as one of the major natural flavorants of cocoa^57,58^. Many other *Geotrichum*/*Dipodascus* species are similarly known for their flavor-enhancing properties^59^.

**Figure 4.**
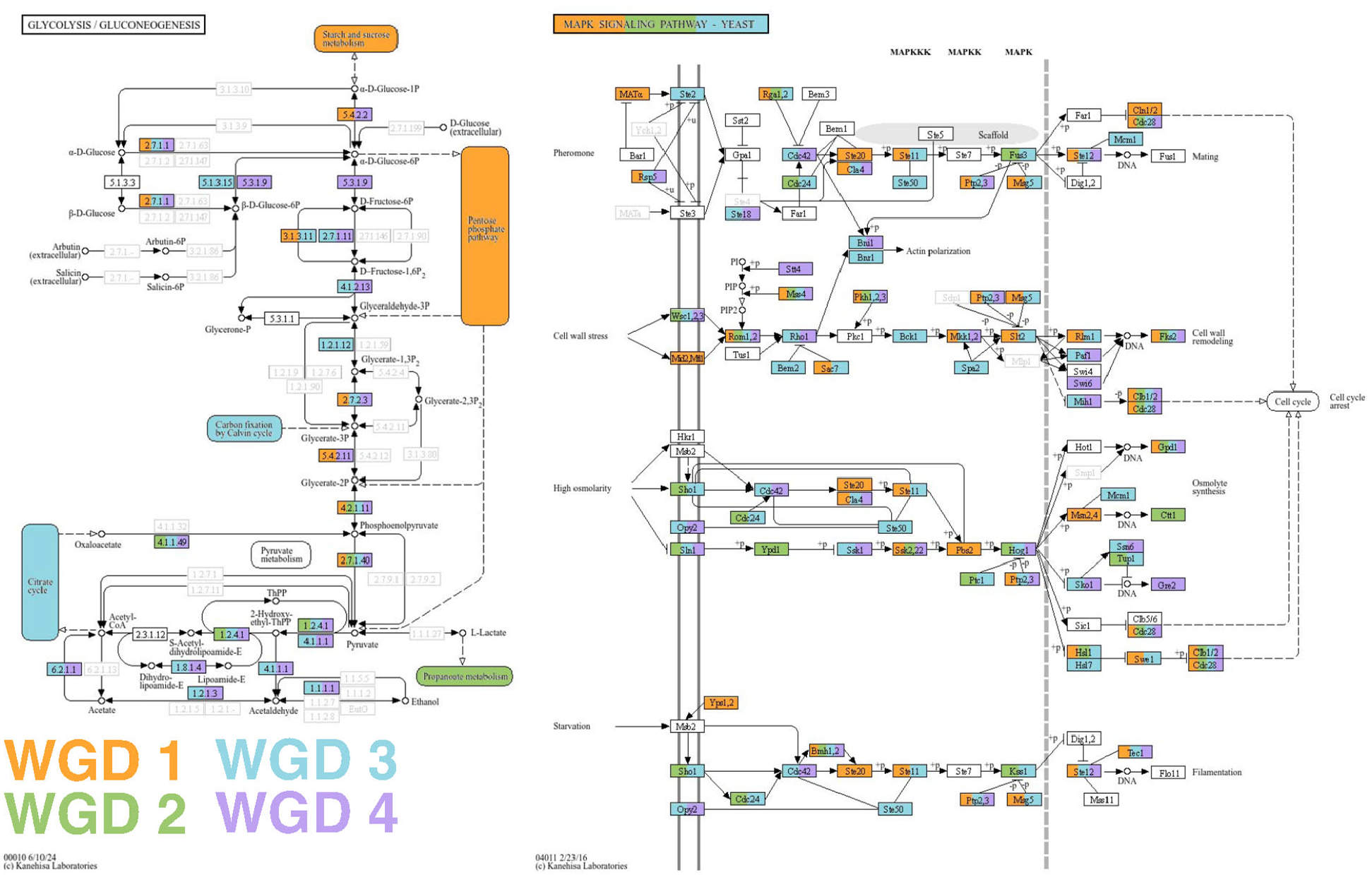
Convergent and divergent impacts of whole genome duplication on metabolic and signaling pathways. Two exemplary KEGG pathways significantly enriched (adjusted p<0.01) among ohnologs from WGD1, WGD2, WGD3, and WGD4 events: glycolysis (left) and the MAPK signaling pathway (right). Genes are colored by which event ohnologs were retained from and pathways are colored by which event they were significantly enriched in. Transparent genes are those that were not identified in any genome in any amount.

As mentioned above, the metabolic effects of WGD1 have been studied previously^48–50,60^. Enzymes and hexose transporters involved in glycolysis have been preferentially retained in post-WGD1 species, in particular those genes whose dosage most greatly increases glycolytic flux^60^. We find all ten of these genes have been retained in duplicate from at least one of the three newly reported WGD events, even those not retained from WGD1. Three gene families, *HXT, ENO*, and *PYK* were retained across all four events. *HXT* is particularly notable, as dosage of these hexose transporter-encoding genes has the single biggest positive impact on glycolytic flux by a wide margin^60^.

It has also been hypothesized that WGD1 coincided with the rise of angiosperms and the increased availability of simple sugars found in fruit and nectar^17^. Our discovery of multiple WGDs reveals that these events are not tied to specific time periods. However, we do find that WGD events are associated with metabolic specialism in yeasts. Post-WGD species metabolize significantly fewer substrates than species that have not undergone WGD (phylogenetic ANOVA p<2.2e-16), a result that persists even if WGD1 taxa are excluded (phylogenetic ANOVA p=1e-3). We hypothesized that additional gene copies in preexisting metabolic pathways produced by WGD events facilitate higher-throughput metabolic output on glucose, reducing evolutionary pressure to metabolize alternative substrates. However, we found that post-WGD species have significantly *reduced* growth rates on the simple sugars mannose (phylogenetic ANOVA p=4.6e-2), and fructose (phylogenetic ANOVA p=1.6e-3), while differences on glucose were insignificant (phylogenetic ANOVA p=0.9). Clearly, more work is required to disentangle the effects of WGD on metabolic rate, growth rate, and niche breadth, an effort we hope will be aided by the discovery of these additional events.

### Expression

While metabolic pathways were differentially enriched across WGD events, other pathways exhibited stronger signatures of convergence. Between KEGG and Gene Ontology annotations, WGD1,2,3 were all significantly enriched with ribosomal proteins (Data S3, Data S4, Fig. S2). In *Saccharomyces* species, the ribosomal protein activator Ifh1 and repressor Crf1 are known ohnologs that diverged from an ancestral generalist regulator following WGD1^61^. Ifh1induces expression under nutrient-rich growth conditions, whereas Crf1 represses expression during periods of stress, providing finer regulatory control during periods of environmental heterogeneity compared to non-WGD1 species^61^. Though these regulators are not found in Dipodascales, regulation of transcription was the third most-enriched gene ontology term in WGD2 ohnologs and the most-enriched in WGD3 ohnologs (Data S3), suggesting similar dynamics may be at play.

### Signaling

Lastly, signaling pathways were also consistent beneficiaries of WGD. The mitogen-activated protein kinase (MAPK), glucagon, and insulin signaling pathways were all significantly enriched (adjusted p<0.01) across WGD1,2,3 (Data S4, Fig. 4, S3). MAPK is an ancient family of kinases that drive phosphorylation cascades, which trigger a variety of cellular mechanisms of *S. cerevisiae* in response to stressful conditions, such as high osmolarity^62^, damage to the cell wall^63^, or starvation^64^ (Fig. 4). While fungi do not naturally produce glucagon or insulin, these general pathways are deeply conserved across eukaryotes^65,66^. These pathways contain protein kinases AMPK and PKA which serve important nutrient sensing functions and trigger cellular responses accordingly^67,68^. Both AMPK and PKA contain duplicate genes from all four WGD events (Fig. S3).

As with yeasts, MAPK proteins in humans exert control over the cell cycle, which render them promising targets for cancer treatments^68,69^. In fact, many pathways significantly enriched (adjusted p<0.01) with ohnologs from WGD1,2,3 are related to human disease, including proteoglycans in cancer, insulin resistance, and COVID-19 (Data S4).

## Conclusion

We report the presence of three previously unknown whole genome duplication events, denoted WGD2, WGD3, and WGD4 in the Dipodascales clade. WGD2 occurred in the most recent common ancestor of the genera *Dipodascus* and *Geotrichum* 201-132mya. WGD3 occurred in the most recent common ancestor of the species *Sap. Suaveolens, M. magnusii,* and *M. tetraspermus* 62-38 mya. WGD4 is specific to *M. magnusii*, occurring 27-0 mya. All these events are strongly supported by both gene/species tree reconciliation and synteny analyses. Despite ∼300 my of evolution separating them, WGD2 and WGD3 share many similar outcomes with the previously known WGD1 occurring in Saccharomycetales, as well as other WGDs across eukaryotes. Genes with many protein-protein interactions, such as those involved with transcription or metabolic/signaling networks, are expected to be more sensitive to dosage effects, and therefore more likely to be retained following WGD^7,70^. Indeed, genes coding for ribosomal proteins and signal transducers have both been preferentially retained following WGD in plants^71^ and animals^72^. The observed extent of convergence suggests that the effects of WGD and other major evolutionary events may be predictable, corroborating recent work in this clade^73^.

Another feature common to WGD-enriched pathways is their role in adaptation under diverse environments. Previous studies in yeasts have shown how these metabolic^60^, ribosomal^61^, and signaling^64,66^ pathways provide heterogenous responses to hostile conditions and to the quantity and quality of available nutrients. The enrichment of these gene families following WGD may explain why modern polyploid *S. cerevisiae* strains are more fit in challenging environments, such as non-optimal carbon sources^74^, human hosts^75^, or brewing vats^76^. This result may further help to address the more widely observed trend of polyploid plant^77^ and animal^78^ species occurring in extreme, rapidly-changing environments.

The nonrandom distribution of retained ohnologs indicates WGD as a potential driver of evolutionary innovation. However, more work is required to identify whether various modes of functional divergence are involved, such as neofunctionalization or escape from adaptive conflict^79^, or if simply increasing dosage of conserved copies is sufficient to precipitate major evolutionary change^60^. Previously thought to be largely absent from fungi^15^, these results underscore the importance of WGD in all eukaryotic kingdoms. We anticipate more fungal WGDs will be discovered as sampling and sequencing continue to improve, and that these events will yield a fuller portrait of eukaryotic genome evolution.

## Methods

### Experimental model and study participant details

*Dipodascus fermentans* PYCC 3480^T^ (NRRL Y-1492) was obtained from the Portuguese Yeast Culture Collection (PYCC) and was routinely grown on solid yeast peptone dextrose (YPD) media at 25ºC. *M. magnusii* (NRRL Y-17563) and *M. tetraspermus* (NRRL Y-7288) were routinely grown on solid yeast peptone dextrose (YPD) media at room temperature (22°C). Liquid cultures were inoculated from a single colony and grown in 25 ml YPD at room temperature in a 125 ml baffled flask shaking at 225 rpm.

## Method details

### D. *fermentans* extraction and assembly

Genomic DNA from overnight grown cultures of *D. fermentans* was obtained using the Quick-DNA Fungal/Bacterial Miniprep Kit from Zymoresearch (cat no. D6005), following the manufacturer’s protocol. Long-read data was obtained using Oxford Nanopore Technology, with a MinION flowcell. For de novo assembly, Canu v2.2^80^ was used with default parameters, only adjusting the genome size flag to 25 m. The resulting contigs were corrected with two rounds of Racon v1.5.0^81^, one with the Nanopore reads and the other with publicly available Illumina reads^82,83^ (SRR16988715). Afterwards, several rounds of Pilon v1.24^84^ were performed using Illumina reads until no changes were seen on the change file. To further increase the contiguity of the assembly, LINKS v1.8.7^85^ was implemented.

### M. *magnusii* and *M. tetraspermus* extraction and assembly

High molecular weight DNA from *M. magnusii* and *M. tetraspermus* was extracted using the Zymo Quick-DNA HMW MagBead Kit (cat no. D6060). The manufacturer’s protocol was followed, but lysis was optimized for non-conventional yeast species. The yeast cells were pelleted by centrifugation at 5000 *x g* and resuspended in 1 ml of 1 M sorbitol with 50 mM dithiothreitol (DTT) and incubated at 30°C for 10 min. The cells were then washed in fresh 1 M sorbitol and resuspended in 200 ul 1 M sorbitol with 5 U of Zymolyase Ultra (Zymo Research, cat no. E1007-2) and incubated at 37°C for 2 hrs. Once cells were spheroplasted, 205 ul phosphate buffered saline, 20 ul of 10% SDS, and 10 ul proteinase K were added, and the spheroplasts were lysed at 55°C for 10 min with occasional inversion. The DNA in the lysate was then bound to the beads following the manufacturer’s protocol. After extraction, the DNA was enriched for high molecular weight DNA using a bead cleanup with a custom buffer (10 mM Tris-HCl, 1 mM EDTA pH 8, 1.6 M NaCl, 11% PEG) as previously described^86^. Genome sequencing was performed by Plasmidsaurus using Oxford Nanopore Technology. Both genomes were assembled with flye v2.9.6^87^ using the `nano-hq` option.

### Gene annotation

To infer gene boundaries for the newly generated genomes of *D. fermentans, M. magnussi*, and *M. tetraspermus* we used funannotate v1.8.16^88^. To do so, each genome was first masked using tantan v40^89^ using the funannotate mask function. Next, each genome was annotated using the funannotate predict function, which is a wrapper function to use multiple gene calling algorithms and creates a consensus set of gene boundaries. Prediction algorithms implemented include Augustus v3.3.2^90^, SNAP v2006-07-28^91^, and GlimmerHMM^92^ each algorithm was trained on gene models predicted using BUSCO v2.0^93^ gene models from the OrthoDB v9^94^ database of near-universally single-copy orthologs from fungi. For Augustus, the ‘optimize_augustus’ argument was used. Additional gene boundaries were predicted by mapping gene annotations from a clustered set of proteins from 332 Saccharomycotina proteomes^82^; clustering was done using CD-HIT v4.8.1^95^ using default settings. The results from each gene prediction algorithm were used to create a consensus set of gene boundaries using EVidenceModeler v1.1.1^96^ with the “repeats2evm” argument. All approaches were given the same weight, except high-quality gene annotations (defined as >90% exon evidence) predicted by Augustus, which were given twice the weight of other algorithms. Gene models less than 50 amino acids in length and putatively transposable elements were subsequently removed. Putative transposable elements were identified using sequence similarity searches conducted using DIAMOND v2.1.8^97^ and the funannotate database of repeat sequences. The resulting gene models were functionally with both KEGG v114.0^54^ and InterPro v106.0^53^ databases. KEGG orthologs were identified through the KofamKOALA^98^ web server. InterPro Gene Ontology annotations were assigned using InterProScan v5.74_105.0^99^. Each gene model was annotated locally using the ‘disable-precalc’ option.

### Gene Tree Inference

Three comparative genomic datasets were used by this study. The first was based on the recently published Y1000+ Project dataset^83^. The full 1,154 genomes of this dataset proved computationally intractable and was subsampled down to 400 using the following procedure: a genome with the shortest terminal branch in the species tree was pruned at random, unless that genome had ≤50 contigs. This process was repeated iteratively until 400 genomes remained in the tree. This sampling strategy maximized phylogenetic breadth and depth while retaining highly-contiguous genomes that could be used for synteny analysis. Gene trees were inferred with OrthoFinder v3.0.1b1^100^ using the Y1000+ species tree as a reference. OrthoFinder identified large spikes in duplication rate within the class Dipodascales, warranting further investigation. Therefore, a second dataset was assembled using all 184 available Dipodascales genomes, as well as a third dataset of 135 Saccharomycetales genomes where WGD1 was known to occur. OrthoFinder was run on both of these datasets as before. A full record of all genomes used in this study can be found at Data S1.

### Quantification and statistical analysis

Synteny analysis was performed using the wgd v2 software package^101^. First, paralogous gene sets were identified within each of 83 Saccharomycotina genome assemblies with ≤50 contigs, in addition to the 2 new genomes sequenced by this study, using the ‘wgd dmd’ command with default parameters. Next, colinear segments (multiplicons) were called with the ‘wgd syn’ command, also under default parameters.

Enrichment analysis was performed using the clusterProfiler v4.10^102^ package in R v4.3.2^103^. The function ‘enrichKEGG’ was used for KEGG pathway enrichment, while the function ‘enricher’ was used for gene ontology enrichment. False discovery rate^104^ was used to control for multiple testing in both cases. Target genes were those that occurred in duplicate across colinear segments within genomes from species predicted to have experienced a given WGD event, against a background of all genes in the genome(s). As *M. magnusii* experienced two rounds of WGD, target genes in this species were further filtered to those estimated to have duplicated prior to the *M. magnusii + Sap. suaveolens* split (for WGD3) and those duplicates specific to *M. magnusii* (for WGD4).

To test whether niche breadth or growth curves were significantly different in post-WGD species, we performed phylogenetic ANOVA using the ‘aov.phylo’ function as implemented in the geiger v2.0.11^105^ package. Niche breadth data was taken from David et al. 2025^73^, and growth curves were obtained from Opulente et al. 2024^83^.

## Acknowledgements

This work was performed using resources contained within the Advanced Computing Center for Research and Education at Vanderbilt University in Nashville, TN. This work was supported by the NSF (grants DBI-2305612 to K.T.D., DEB-2110403 to C.T.H., and DEB-2110404 to A.R.) and the NIH (R35GM151348 to M.P.). This work was partially supported by FCT—Fundação para a Ciência e a Tecnologia, I.P. (FCT/MCTES; https://www.fct.pt/) in the scope of projects UIDP/04378/2020, UIDB/04378/2020, LA/P/0140/2020 and grant PTDC/BIA-EVL/0604/2021 (to C.G.). Research in the Hittinger Lab is also supported by the United States Department of Agriculture National Institute of Food and Agriculture (Hatch Project 7005101), in part by the Department of Energy (DOE) Great Lakes Bioenergy Research Center (DOE Biological and Environmental Research Office of Science DE–SC0018409). Research in the Rokas Lab is also supported by the NIH/National Institute of Allergy and Infectious Diseases (R01 AI153356). J.L.S. is a Howard Hughes Medical Institute Awardee of the Life Sciences Research Foundation.

## Conflicts of Interest

J.L.S. is an advisor to ForensisGroup Inc. J.L.S. is a scientific consultant to FutureHouse Inc. A.R. is a scientific consultant for LifeMine Therapeutics, Inc. All other authors declare no conflict of interest.

