## Supplementary figures and images for "Discovery of additional ancient genome duplications in yeasts"

### Supplementary Figures

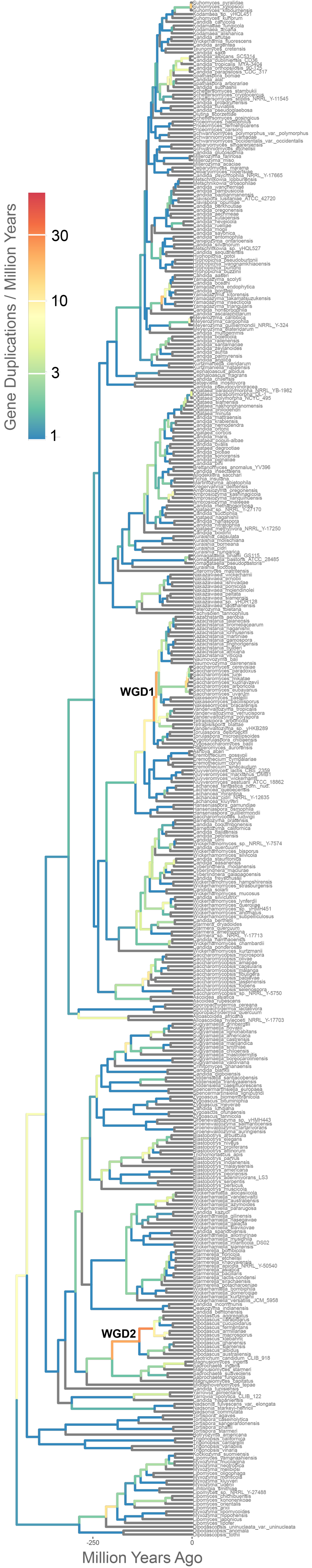

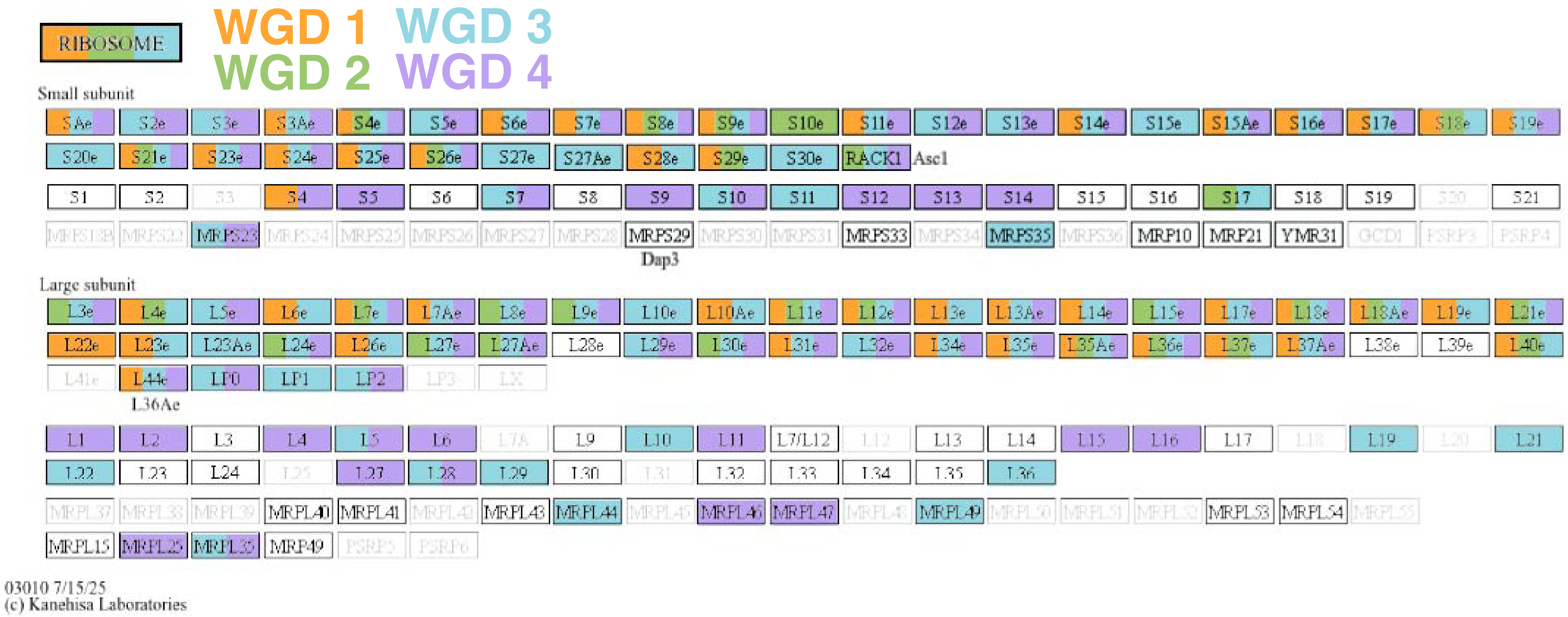

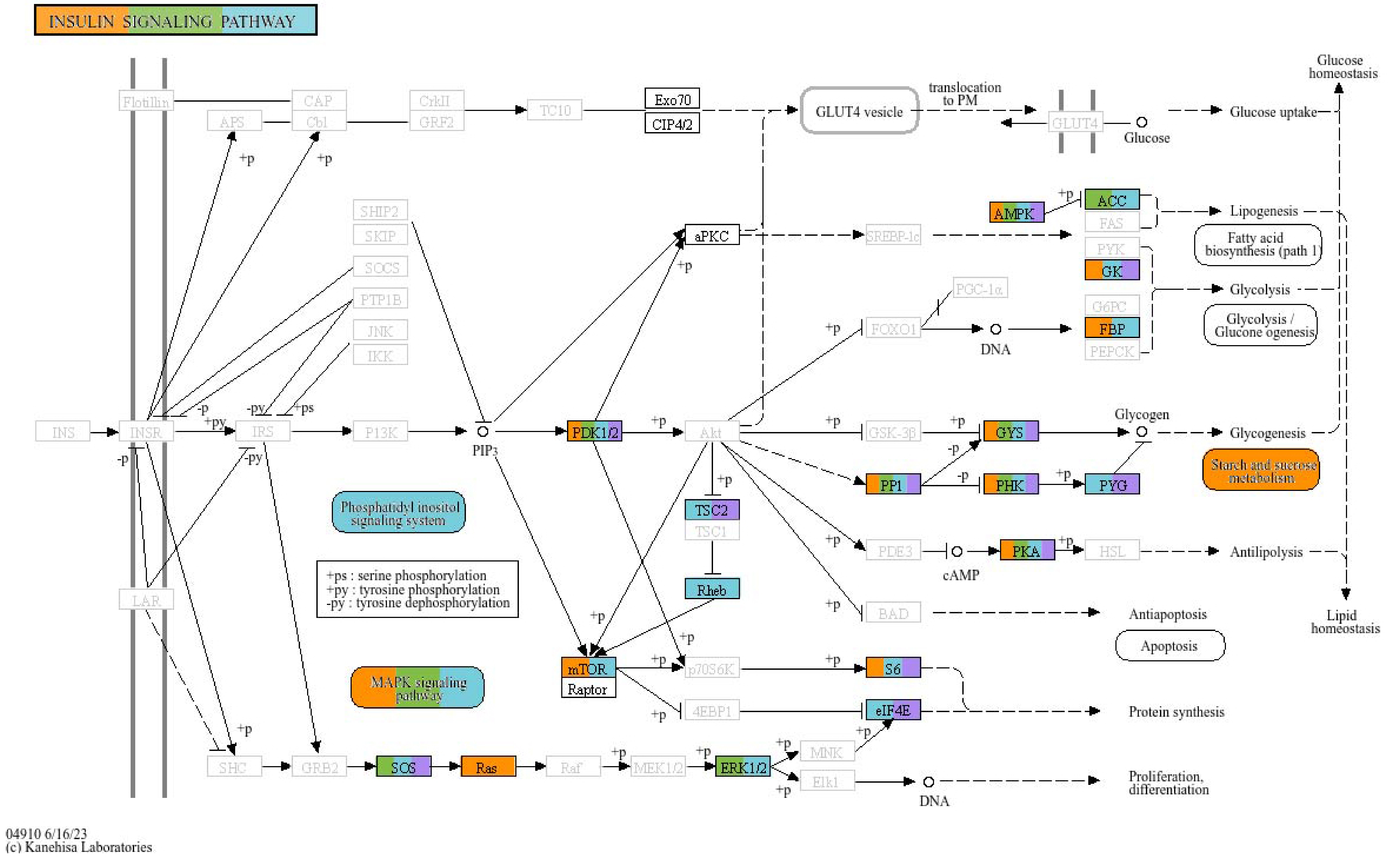

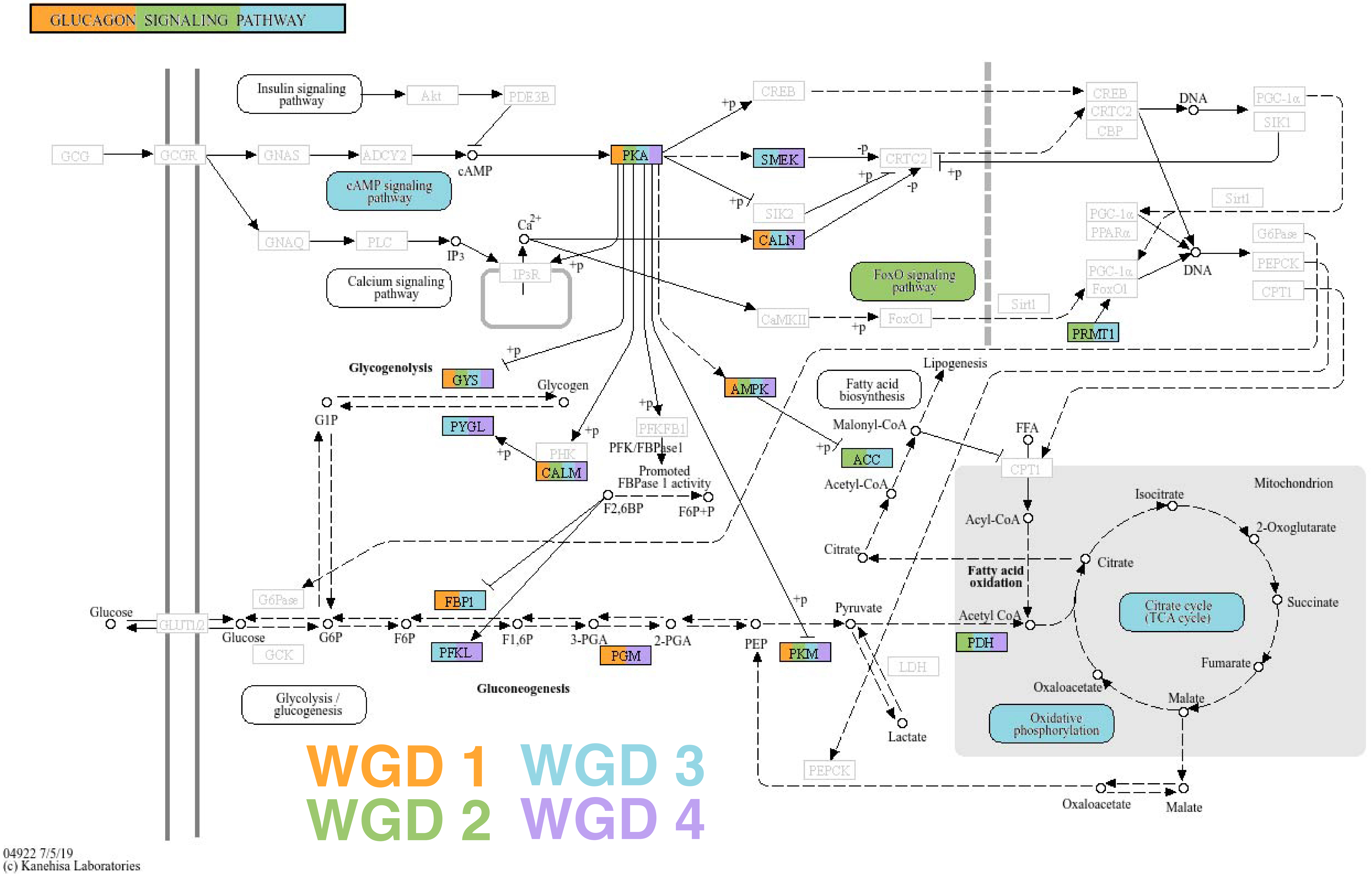
